# Langerhans cells immunocompetency is critical for IDO1-dependent ability to induce tolerogenic T cells

**DOI:** 10.1101/2019.12.20.884130

**Authors:** James Davies, Sofia Sirvent, Andres F. Vallejo, Kalum Clayton, Gemma Porter, Patrick Stumpf, Jonathan West, Michael Ardern-Jones, Harinder Singh, Ben MacArthur, Marta E Polak

## Abstract

Human epidermal Langerhans cells (LCs) can coordinate both immunogenic and tolerogenic immune responses, creating an attractive opportunity for immunomodulation strategies. To investigate transcriptional determinants of human primary LC tolerance we applied single cells RNA-sequencing combined with transcriptional network modelling and functional analysis. Unsupervised clustering of single cell transcriptomes revealed that steady-state LCs exist in immature and immunocompetent states, and become fully immunocompetent on migration. Interestingly, LC migration, which has been shown to result in upregulation of the transcription factor IRF4, led in parallel to increased expression of a tolerogenic gene module including *IDO1*, *LGALS1*, *LAMTOR1* and *IL10RA,* which translated to efficient induction of regulatory T cells in co-culture assays by immunocompetent LCs. Using protein expression analysis and perturbation with inhibitors, we confirmed the role of IDO1 as a mediator of LC tolerogenic responses induced during LC migration. Computational analysis of regulons and Partial Information Decomposition analyses identified *IRF4* as a key driver for LC tolerogenic programmes. The predicted IRF4-regulated genes were confirmed by analysis of CRISPR-Cas9 edited LCs. These findings suggest that efficient priming of tolerogenic responses by LCs requires upregulation of a migration-coupled maturation program which is superimposed with a tolerance-inducing genomic module.

## Introduction

Langerhans cells (LCs) reside in the epidermis as a dense network of immune system sentinels, capable of initiating potent immune responses to cutaneous pathogens and neoplastic cells^1,2^. As a first line of the cutaneous immune defence system, LCs are uniquely specialised at sensing the environment and extend dendrites through inter-cellular tight junctions to gain access to the outermost part of the skin, the stratum corneum, so that rapid responses can be initiated if a dangerous pathogen is encountered^3^. We and others have shown that LCs are highly capable of priming and augmenting CD4 T cell responses and can induce CD8 T cell activation via antigen cross presentation more effectively than CD11c+, CD141+, CD141- and CD14+ dermal DCs^4,5,6^. However, in the context of infection, LCs can be surprisingly inefficient; LCs fail to induce cytotoxic T cell in response to herpes simplex 1 virus and in the context of *Leishmania major* infection, ablation of LCs reduced the number of activated Treg cells and aided clearance of the disease, therefore questioning the role of LC in immunogenic responses to pathogenic stimuli^7,8^.

In contrast, during steady-state (non-dangerous) conditions, LCs selectively induce the activation and proliferation of skin-resident regulatory T cells^9,10^ that prevent unwanted immune-mediated reactions. This key role of LCs in the maintenance of cutaneous and systemic homeostasis has been confirmed in many experimental systems. Using a mouse model for investigating LC tolerance, antigen processing and presentation of the keratinocyte protein desmoglein (Dsg3) resulted in efficient regulatory T cell (Treg) induction, whilst ablation of LCs led to increased autoimmunity^11^. LC migration from the epidermis is constantly observed during steady-state and transport of self-antigen derived from melanin to skin draining lymph nodes results in no abnormal inflammatory disease, nor does the rate of transport change during induced inflammatory conditions^12,13,14^. In a mouse model of autoimmune encephalomyelitis, migratory skin LCs and not resident lymph DCs are required for the induction of Foxp3+ Tregs^15^. Epicutaneous delivery of OVA to OVA-sensitised mice and peanut protein to peanut-sensitised mice results in uptake and processing by skin DC and LC, but repeat exposure results in fewer inflammatory responses and an increase in Treg cells^16,17^. Similarily, LCs mediate murine tolerance during sensitisation by the hapten DNTB and is reliant on the presence of Tregs^18^.

While LC-induced tolerance appears to be key to cutaneous immune homeostasis, and understanding of molecular processes underpinning this can open opportunities for targeted vaccine and immunosuppressive therapeutics, currently little is known about what biological pathways LC use for directing tolerogenic T cell immune responses. Even though some comparisons can be drawn from analysis of other tolerogenic dendritic cell subsets, LC transcriptional programming is distinct from both dendritic cells and macrophages^19,20,5,21^. Thus, in-depth analysis of human primary LCs is necessary to expand understanding of their ability to induce tolerogenic responses. Here, we used single cell RNA-seq and *in vitro* experimentation to advance understanding of LC heterogeneity and immune activation at both the steady-state and after migration, revealing important insights into how LCs mediate tolerogenic T cell responses at the epidermis.

## Methods

### Human LC and PBMC isolation

Human blood and skin mastectomy and abdominoplasty samples were collected with written consent from donors with approval by the South East Coast - Brighton & Sussex Research Ethics Committee in adherence to Helsinki Guidelines [ethical approvals: REC approval: 16/LO/0999). Fat and lower dermis was cut away and discarded before dispase (2 U/ml, Gibco, UK, 20h, +4°C) digestion. For steady-state LC extraction, epidermal sheets were digested in Liberase Tm (13 U/ml, Roche, UK, 2h, +37°C,) and enriched using density gradient centrifugation (Optiprep 1:4.2, Axis Shield, Norway). Migrated LCs were extracted from epidermal explant sheets cultured in media (RPMI, Gibco, UK, 5%FBS, Invitrogen, UK, 100 IU/ml penicillin and 100 mg/ml streptomycin, Sigma, UK) for 48 hours, with or without dexamethasone (1μM, Hameln, UK). Dexamethasone stimulated migrated LC were washed with media prior to use in assays. Steady state and migrated LC were processed through fluorescence-activated cell sorting (FACS) and Drop-seq or cryopreserved in 90% FBS (Gibco, UK), 10% DMSO (Sigma, UK). PBMCs were extracted from human blood using lymphoprep (Stemcell, UK) density gradient separation. Naïve T cells were purified using the Naïve CD4+ T cell isolation kit (Miltenyi Biotec, UK). TRMs were extracted from epidermal sheets after 48 hour migration, followed by density gradient separation (Optiprep 1:3).

### Flow cytometry/ FACS

Antibodies used for cell staining were pre-titrated and used at optimal concentrations. A FACS Aria flow cytometer (Becton Dickinson, USA) and FlowJo software was used for analysis. For FACS purification LCs were stained for CD207 (anti-CD207 PeVio700), CD1a (anti-CD1a VioBlue) and HLA-DR (anti-HLA-DR Viogreen, Miltenyi Biotech, UK). For T cell staining, antibodies anti-CD3 PerCP, anti-CD4 Viogreen, anti-CD127 Pe (Miltenyi Biotech, UK) and anti-CD25 PeCy7 (Invitrogen, UK) were used for surface staining. Anti-FOXP3 FITC (eBiosciences, UK), anti-IL-10 PE (Miltenyi, UK) and anti-IDO1 AlexaFluor647 (Biolegend, UK) antibodies were used for intranuclear and intracellular staining.

### Co-culture, suppression and inhibition assays

For co-culture assays, purified LC and naïve CD4+ T cells or TRMs were co-cultured in human serum supplemented media (RPMI, Gibco, UK, 10% human serum, Sigma, UK, 100 IU/ml penicillin and 100 mg/ml streptomycin, Sigma, UK) at a 1:50 ratio for 5-days at 37°C. For intranuclear FOXP3 staining T cells were permeabilised using the FOXP3/Transcription Factor Staining Buffer Set (eBiosciences, UK) following the manufacturers protocol, after cell surface marker staining. For IL-10 intracellular staining, T cells were stimulated with cell stimulation cocktail (eBioscience, UK) for 6 hours and Golgi plug (eBioscience, UK) for 5 hours, prior to intracellular staining using Permeabilizing Solution 2 (BD Biosciencies, UK). IDO1 intracellular staining of LCs was performed using Intracellular Fixation & Permeabilization Buffer Set (eBioscience, UK), following kit protocol. IDO1 inhibition experiments were performed using NLG-919 (10μM, Cambridge Bioscience UK) and epacadostat (EPAC, 1μM, Cambridge Bioscience UK) in media during migrated LC and naïve CD4+ T cell co-cultures. Proliferation assays were set up through combining FACS-purified CD3+CD4+CD127-CD25+ T cells induced after 5-day naïve CD4+ T cells and FACS-purified LC co-cultures, with autologous CFSE labelled PBMCs. PBMCs were labelled with CFSE using the CellTrace™ CFSE Cell Proliferation Kit (Invitrogen, UK), with ice cold PBS, 0.5% BSA replacing PBS and ice cold media replacing pre-warmed media as described in the protocol.

### Drop-seq

After FACS purification, single LCs were co-encapsulated with primer coated barcoded Bead SeqB (Chemgenes, USA) within 1 nL droplets (Drop-seq^22^). Drop-seq microfluidic devices according to the design of Macosko *et al* were fabricated by soft lithography, oxygen plasma bonded to glass and functionalised with fluorinated silane (1% (v/v) trichloro(1H,1H,2H, 2H-perfluorooctyl)silane in HFE-7500 carrier oil). Open instrumentation syringe pumps and microscopes (see dropletkitchen.github.io) were used to generate and observe droplets, using conditions and concentrations according to the Drop-seq protocol^22^, 700 steady-state LC and ~300 migrated LC from mastectomy skin were converted into ‘STAMPs’ for PCR library amplification (High Sensitivity DNA Assay, Agilent Bioanalyser) and tagmentation (Nextera XT, Illumina, UK). Sequencing of libraries was executed using NextSeq on a paired end run (1.5×10E5 reads for maximal coverage) at the Wessex Investigational Sciences Hub laboratory, University of Southampton.

### Transcriptomic data analysis

The Drop-seq protocol from the McCarrol lab^22^ was followed for converting sequencer output into gene expression data. The bcl2fastq tool from Illumina was used to demultiplex files, remove UMIs from reads and deduce captured transcript reads. Reads were then aligned to human hg19 reference genome using STAR. Analyses of steady-state LCs alone was performed in an R environment using SCnorm normalisation ^23^, universal manifold approximation and projection (UMAP) dimensionality reduction analysis (Scater^24^, singlecellTK^25^) and hierarchical clustering (clust=ward.D2, dist = canberra)^26^. Differentially expressed genes (DEGs) between cell clusters were identified using Limma^27^ (FDR corrected p-value<0.05, logFC>1). Comparative analyses of steady-state and migrated was performed using the python-based Scanpy pipeline (version 1.5.0) ^28^. High quality barcodes, discriminated from background RNA barcodes, were selected based on the overall UMI distribution using EmptyDrops ^29^. Low quality cells, with a high fraction of counts from mitochondrial genes (20% or more) indicating stressed or dying cells were removed. In addition, genes with expression detected in <10 cells were excluded. Datasets were normalised using Scran, using rpy2 within python ^30^. Highly variable genes (top 2000) were selected using distribution criteria: min_mean=0, max_mean=4, min_disp=0.1. A single-cell neighbourhood graph was computed on the first principal components that sufficiently explain the variation in the data using 10 nearest neighbours. Uniform Manifold Approximation and Projection (UMAP) was performed for dimensionality reduction. Leiden algorithm (Traag, Waltman and van Eck, 2019, PMID: 30914743) was used to identify clusters within cell populations (Leiden r = 0.5, n_pcs=30). Differentially expressed genes (DEGs) between cell clusters were identified using T-test (FDR corrected p-value<0.01, logFC>1). Gene ontology analysis was performed using Toppgene (FDR corrected p-value<0.05), describing biological pathways associated with gene lists. Tolerogenic gene signature 1 (tol 1) was curated from published studies and literature exploring genes associated with DC or macrophage tolerogenic function. Single cell gene signature enrichment analyses of tol 1 was performed using Gene Set Variation Analysis (GSVA) ^31^. Regulatory network inference analysis was performed using single-cell regulatory network inference and clustering (SCENIC) within python (Aibar et al., 2017). Public datasets from GEO used for analysis included a microarray dataset containing dexamethasone and vitamin D3 stimulated MoDC (TolMoDC) with unstimulated MoDC (GSE52894) and a microarray dataset containing trypsinised steady-state LC with unstimulated MoDC (GSE23618) Normalised count matrices were downloaded from GEO before Limma DEG analysis in R. DEGs upregulated in LCs and TolMoDCs compared to unstimulated MoDCs from each respective dataset were anlaysed, with unstimulated MoDC used as reference for comparison.

#### Directional PIDC

Notebooks from Chan et al were adapted for the analysis and run using Julia V 1.0.5 in Jupyter Notebook. SCRAN-normalised data for migrated LCs including genes from the tol1 signatures and selected transcription factors was used for network inference using PIDC algorithm. Edge weights were exported, and sorted to include only transcription factors as targets. Hierarchical network was visualised using yED.

## Results

### Steady state LCs exist in a spectrum of immune activation from immaturity to immunocompetency

LCs in steady-state healthy skin have been shown to expand skin resident memory Tregs which are important for mediating immune homeostasis and preventing unwarranted inflammatory responses^9^. However, when compared to dexamethasone and vitamin D3 stimulated model tolerogenic dendritic cells (TolMoDC), trypsinised steady-state LC display unique upregulated biological pathways with no crossover of differentially expressed genes (DEGs) when compared to unstimulated TolMoDC (FigureS1A, Figure S1B, Supplementary table 1). To explore the gene expression profiles underlying healthy LC tolerogenic function and to evaluate population heterogeneity *in situ* we performed single cell RNAseq on “steady-state LC” dissociated from healthy skin using the dispase/liberase protocol, as published previously^32,33^. UMAP dimensionality reduction analysis of 607 steady-state LCs revealed heterogeneity was present amongst the population, with LCs transitioning from one state to another (Figure 1A). Using hierarchical clustering, steady-state LCs were grouped into two defined clusters (S1 and S2) which separated the population along the transitional route (Figure 1B). Gene expression comparison after grouping genes into intervals of increasing expression level, revealed S2 LCs to be more transcriptionally active than S1 (Figure 1C). Whilst the most highly expressed genes (>10 normalised expression) in both sub populations were associated with antigen processing and presentation, the number of genes and range of expression was highest at all expression intervals in S2 LCs. Differentially expressed gene (DEG) analysis identified 21 upregulated genes (*CD74, HLA-DRA, HLA-DRB1, B2M) in* S2 LCs compared to S1 LCs, although no DEGs were identified in S1 compared to S2 (Supplementary table 2). Whilst no specific gene ontologies were associated with tolerogenic pathways, analysis revealed associations with MHC II antigen presentation, T cell co-stimulation and response to cytokines (Figure 1D). Overall, steady-state LCs appear to exist in a spectrum of activation from lowly activated immature LC (S1) to more highly activated immunocompetent LC (S2), which likely influences their potential for coordinating T cell responses.

**Figure 1.**
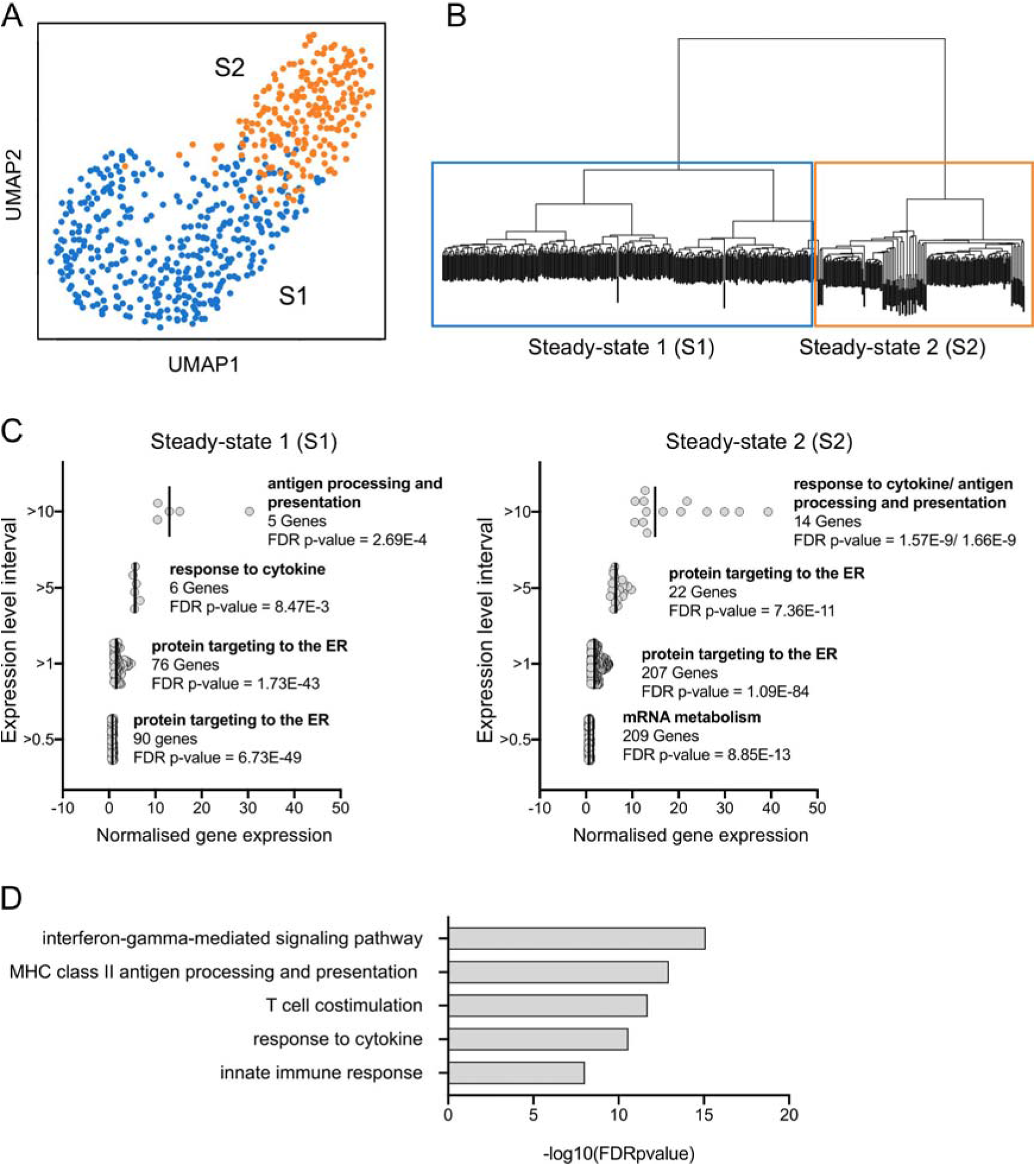
Steady state LCs exist in a spectrum of immune activation from immaturity to immunocompetency. A. UMAP plot of 607 steady-state LCs (Scater, R) from 1769 genes following filtering (mitochondrial genes <20%) and SCnorm normalisation. B. Hierarchical clustering (clust=ward.D2, dist=canberra) of steady-state LCs defining the division of the population into two sub clusters, steady-state 1 (S1) and steady-state 2 (S2). C. S1 and S2 normalised gene expression values were grouped into expression level intervals (y-axis). The number of genes included in each interval and the top associated biological processes identified using ToppGene gene ontology analysis are displayed with significance values (FDR corrected p-value). D. Gene ontology analysis using ToppGene of 21 DEGs upregulated in S2 LC compared to S1 LC. -log(10) FDR corrected FDR corrected p-values are displayed.

### Steady state immunocompetent LCs are superior at inducing FOXP3+ Treg

As activation status defined LC subpopulations in the steady-state, the expression of classical DC activation markers and LC markers amongst the population was interrogated (Figure 2A, Figure S2A). Whilst *CD83* expression appeared homogenous, *CD40* was lowly detected in all steady-state LC. High HLA-DRA expression was detected in LCs from both clusters but *CD86* was predominantly expressed in S2 LC only. Following confirmation of the differential expression at the protein level using flow cytometry, CD86 was selected as a marker for distinguishing the two populations: immunocompetent and immature LCs (Figure 2B). To investigate the immune potential of these two steady-state LC populations, CD86High and CD86Low expressing LCs were isolated using FACS (Figure 2B) and co-cultured with CD4+ naïve T cells for 5-days after which the expansion of CD25+FOXP3+ Tregs was quantified (Figure 2C, Figure S2B). Suprisingly, CD86Low immature LCs did not increase the number of CD4+CD25+FOXP3+ regulatory T cells compared to control. In contrast, CD86High immunocompetent LC significantly expanded the number of CD25+FOXP3+ Tregs compared to control (p=0.0143) and CD86Low LC (p=0.0129, n=3 independent skin donors), revealing that the state of immunocompetence associates with LC ability to promote T cell-mediated immune tolerance.

**Figure 2.**
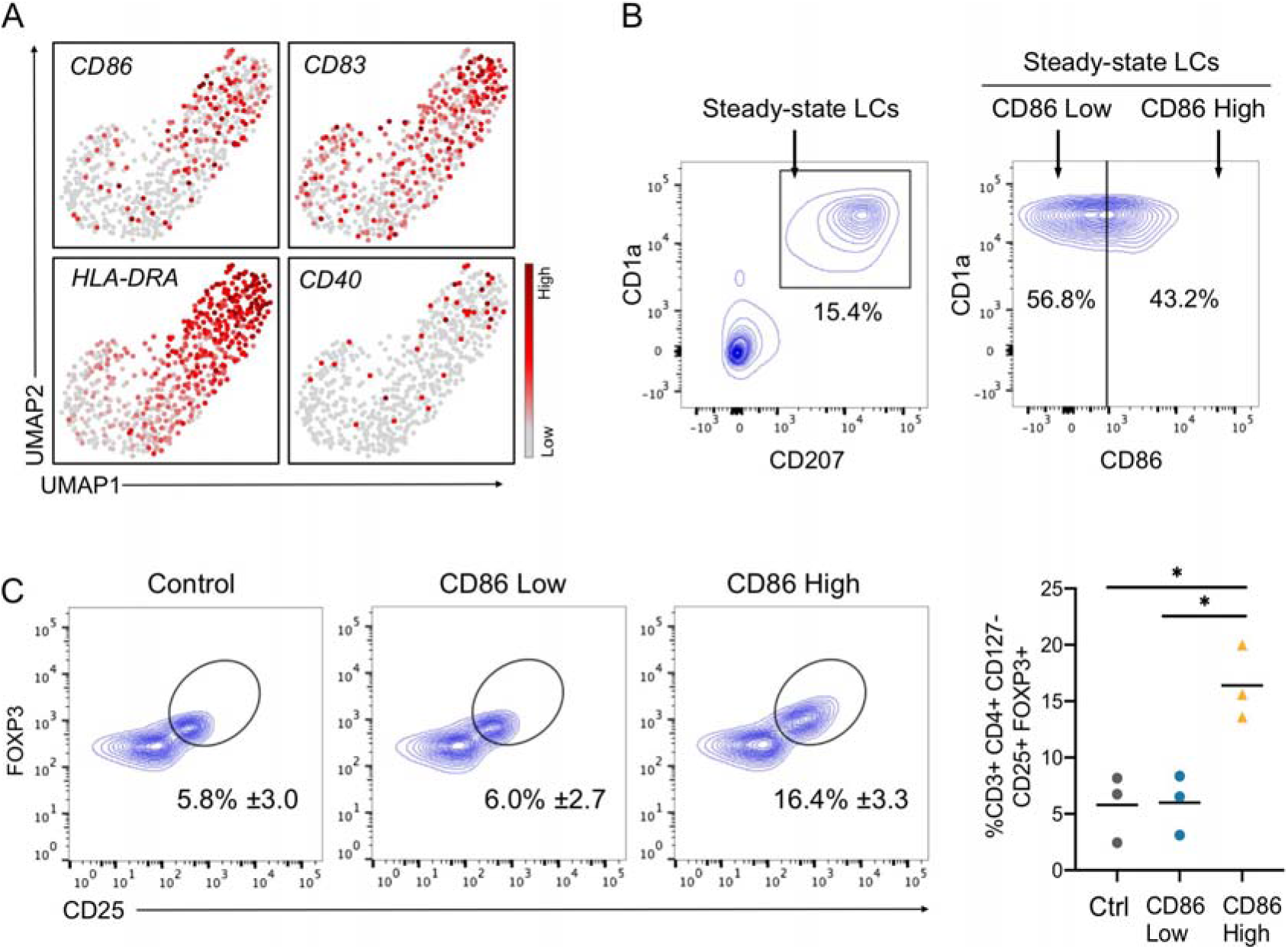
Steady state immunocompetent LCs are superior at inducing FOXP3+ Tregs. A. UMAP plots displaying markers: *CD86, CD83*, *HLA-DRA* and *CD40* expression amongst the steady-state LC population displaying low (grey) to high (dark red) SCnorm normalised expression. B. Flow cytometry assessment of steady-state LCs identified as CD207/CD1a high cells. LC populations were separated into CD86Low and CD86High by FACS. Representative example from n=3 independent LC donors. C. Flow cytometry assessment of CD4+ naive T cells after 5-day co-culture with either CD86Low or CD86High steady-state LC. 5-day cultures of CD4+ naïve T cells alone were used as control. Tregs were identified as CD3+CD4+CD127-CD25+FOXP3+ cells. n=3 independent LC donor paired experiments. *p<0.05.

### Migration enhances tolerogenic abilities of immunocompetent LCs

To explore further the link between LC activation status and tolerance induction, we sought to investigate the effect of i*n vitro* migration from epidermal sheets on LCs tolerogenic characteristics and their molecular profile. We and others have previously shown that LCs, which have migrated out of the epidermis are more immune activated and primed to mediating T cell responses^6,32,33^. Therefore, we sought to compare the capability of both steady-state LC and migrated LC to prime naïve CD4 T cells towards a tolerogenic phenotype. The expansion of CD25+FOXP3+ Tregs was measured after 5-day co-culture of naïve CD4+ T cells with LCs extracted at the steady-state via 2 hour enzymatic digestion or through migration from epidermal explants (Figure 3A). Steady-state and migrated LC both significantly amplified the percentage of CD25+FOXP3+ Tregs compared to CD4-only control (n=6 independent skin donors, steady-state LCs p=0.0101, migrated LCs p=<0.0001). However LCs ability to amplify Tregs was significantly augmented after migration, with increased percentages of CD25+FOXP3+ Tregs induced compared to steady-state LCs (Figure 3A, p=<0.0001). When co-cultured with antiCD3/CD28-stimulated PBMCs, Tregs expanded with migratory LCs potently inhibited activated CD4 and CD8 T cell proliferation (Figure 3B, C, respectively and Figure S3A, CD4 1:1 p=0.0088, CD4 1:3 p=0.0277, CD8 1:1 p=0.0007, CD8 1:3 p=0.0111, n=5 from 3 independent LC donors). Similarily, migrated LCs efficiently expanded autologous epidermal tissue-resident memory T cells (TRMs) isolated from healthy epidermal tissue (Figure 3D, Figure S3B). Co-culture of migrated LCs with TRMs significantly increased the number of CD25+FOXP3+ Tregs compared to steady-state control (n=5 steady-state LC independent skin donors, n=4 migrated LC independent skin donors, p=0.0025). Furthermore, co-culture of migrated LCs with resident memory T cells also drove expansion of IL-10 producing CD4+ T cells, highlighting the tolerogenic capabilities of migrated LCs (Figure 3E, n=8 independent skin donors, p=0.0451).

**Figure 3.**
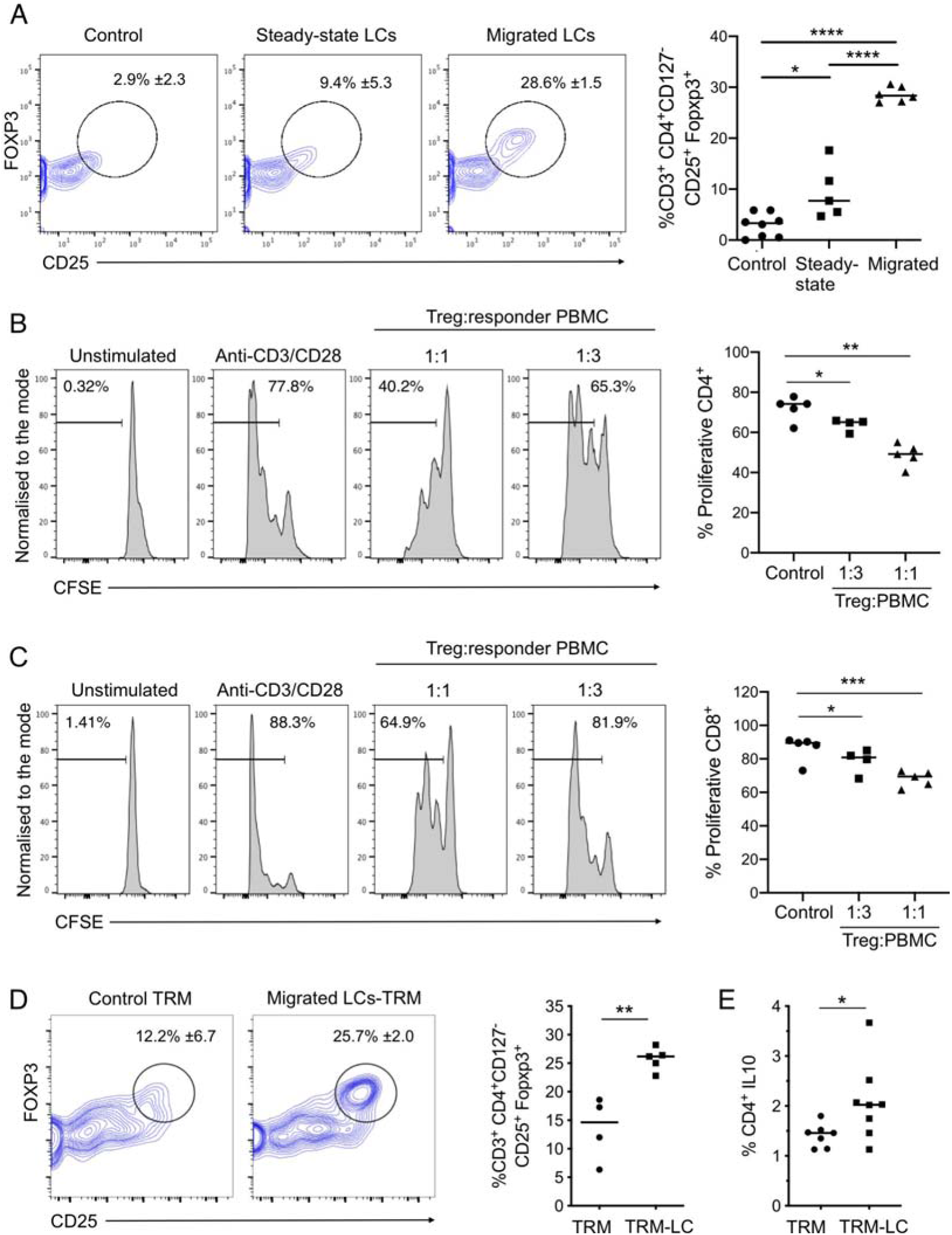
Migration enhances tolerogenic abilities of immunocompetent LCs. A. Flow cytometry assessment of the percentage of Tregs induced after 5-day co-culture of steady-state LC and migrated LC with CD4+ naive T cells. 5-day cultures of CD4+ naïve T cells alone were used as control. Tregs were identified as CD3+CD4+CD127-CD25+FOXP3+ cells. n=8 control, 5 steady-state and 6 migrated independent LC donors. *p<0.05, **p<0.01, ***p<0.001. B. CFSE labelled PBMCs gated on CD4+ T cells proliferation measurements after 3-day co-culture with autologous purified CD3+CD4+CD127-CD25+ Tregs. The percentage of proliferating CD4+ cells stimulated with plate bound anti-CD3 and soluble anti-CD28 is displayed at ratios of 1:1 and 1:3 Treg:PBMC (n=5 from 3 independent LC donors). *p<0.05, **p<0.01. C. CFSE labelled PBMCs, gated on CD8+ T cells, proliferation measurements after 3-day co-culture with autologous purified CD3+CD4+CD127-CD25+ Tregs. The percentage of proliferating CD8+ cells stimulated with plate bound anti-CD3 and soluble anti-CD28 is displayed at ratios of 1:1 and 1:3 Treg:PBMC (n=5 from 3 independent LC donors). *p<0.05, ***p<0.001. D. Flow cytometry assessment of the percentage of Tregs induced after 5-day co-culture of migrated LC with autologous T resident memory cells (TRMs) extracted from human epidermis. 5-day cultures of TRMs alone were used as control. Tregs were identified as CD3+CD4+CD127-CD25+FOXP3+ cells. n=5 independent LC donors. **p<0.01. E. Percentage of IL-10 producing CD4+ cells after co-culture of TRMs with or without migrated LC. n=8. *p<0.05.

### Migration from the epidermis enhances LCs tolerogenic programming

The induction of tolerance by LC is believed to be critical for maintaining homeostasis at the skin ^9,20,34^. To investigate transcriptional programmes encoding ability of LCs to induce immunotolerance, we first assembled a tolerogenic DC gene signature panel, based on literature review (tol 1, Supplementary Table 3). We confirmed that the signature was significantly enriched in LCs, either at the steady-state or migrated, with LCs expressing 30/64 genes of the tol1 signature (Figure 4A,B, Supplementary Figure 4A,B). Surprisingly, the signature had significantly higher enrichment in migrated LCs (tol1= 1.26E-14), compared to steady state LCs, with 11/30 genes significantly upregulated in migrated LC (p<0.05, logFC>1, Figure 4B,C, Supplementary Table 4).

**Figure 4.**
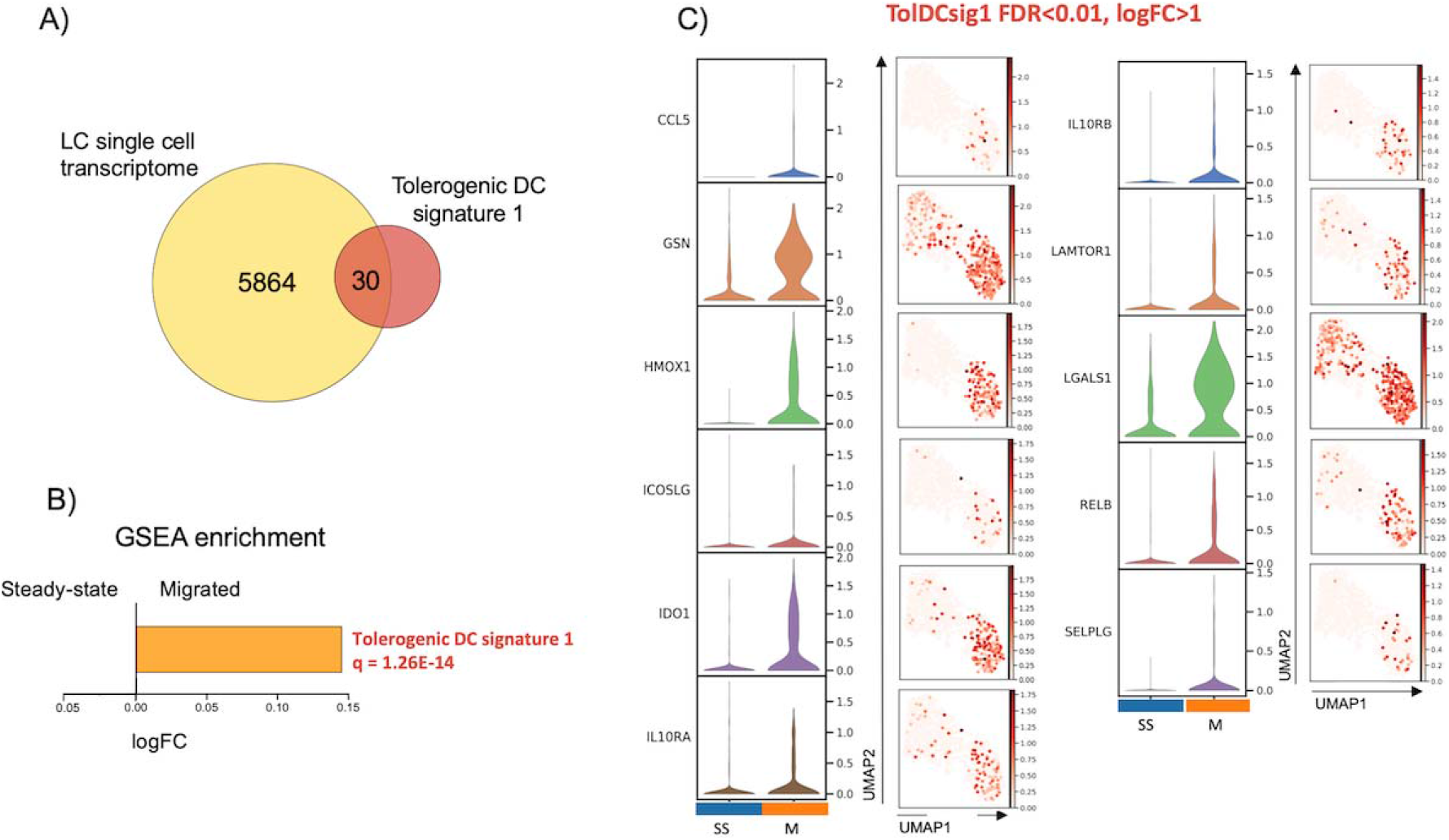
Migration from the epidermis enhances LCs tolerogenic programming. A. Venn diagram displaying the number of genes from tolerogenic gene signature 1 (tol 1), curated from literature exploring genes associated with DC or macrophage tolerogenic function, within the whole LC single cell dataset. B. Gene Set Variation Analysis (GSVA) displaying enrichment of tol 1 in the LC populations. FDR corrected p-values and logFC are displayed. C. Violin plots and UMAP marker plots displaying the expression of genes within tol 1 amongst steady-state and migrated LCs (FDR corrected p-values <0.01, logFC>1).

### LC-induced tolerance is mediated by IDO1 and can be enhanced by immunotherapeutic intervention

Among all potential mediators of immunotolerance in immunocompetent LCs, *IDO1* was the most extensively expressed in migrated LC, and homogenous in this population. Consistent with the single cell RNA-seq data, the level of IDO1 protein expression was considerably and significantly higher in migrated LCs compared to steady-state LC (Figure 5A, Figure S5A, n=5 independent skin donors, p=0.0002), indicating that this molecule can be critical to migrated LC tolerogenic function. Blocking of IDO1 signalling with NLG-919, an immune checkpoint inhibitor, significantly impaired LCs ability to expand tolerogenic T cells (Figure 5B, p=0.0354). Interestingly, interference with IDO1 using epacadostat (EPAC), another selective inhibitor of tryptophan catabolism was less potent (Figure 5B, p=0.0583). LCs ability to prime and expand tolerogenic T cells creates an exciting opportunity for therapeutic interventions. Since steady-state LCs exist in a spectrum of immunocompetence, with a subpopulation of LCs already poised for tolerance induction, we hypothesised that *in situ* treatment can further potentiate their tolerogenic behaviour upon migration. To test this, we treated LCs with dexamethasone during migration from the epidermis. Indeed, dexamethasone migrated LCs were significantly more potent in expanding CD25+FOXP3+ Tregs (Figure 5C, n=4 independent skin donors, p=0.0271) in comparison to their untreated migrated counterparts. Additionally, CD4+ T cells expanded by migratory DexLCs (n=5 independent skin donors, p=0.0061) produced more IL10 than untreated migrated LC (p=0.028), consistent with their tolerogenic phenotype (Figure 5D). Importantly, the presence of dexamethasone during LC migration further increased the expression of IDO1 protein and therefore enhanced the tolerogenic LC phenotype (n=4 independent skin donors, p=0.0142), supporting the importance of IDO1 for LC tolerogenic function (Figure 5A, Figure S5A).

**Figure 5.**
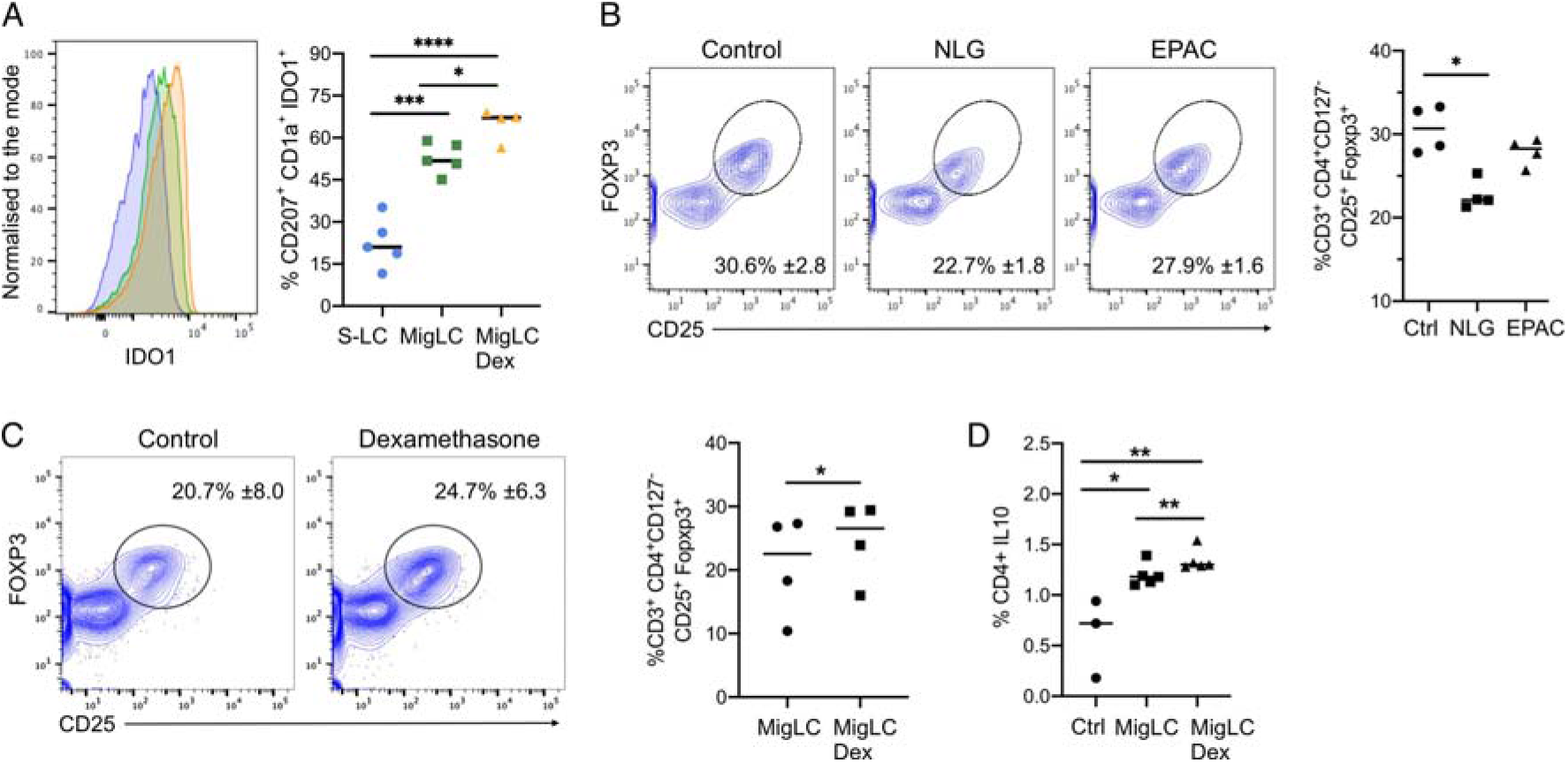
LC-induced tolerance is mediated by IDO1 and can be enhanced by immunotherapeutic intervention. A. Flow cytometry assessment of the percentage of IDO1 expression in steady-state LC and migrated LC extracted by 48 hour culture of epidermal sheets with and without 1M dexamethasone. n=5 steady-state and migrated independent LCs, n=4 migrated dexamethasone independent LCs. *p<0.05, ***p<0.001, ****p<0.0001. B. Flow cytometry analysis of the percentage of Tregs induced after 5-day co-culture of migrated LC with CD4+ naïve T cells in the presence of IDO1 inhibitors NLG-919 (NLG) and epacadostat (EPAC). 5-day cultures of CD4+ naïve T cells alone were used as control. Tregs were identified as CD3+CD4+CD127-CD25+FOXP3+ cells. n=4 independent LC donors. *p<0.05. C. Flow cytometry assessment of the percentage of Tregs induced after 5-day co-culture of migrated LC with and without dexamethasone stimulation, with CD4+ naive T cells. Tregs were identified as CD3+CD4+CD127-CD25+FOXP3+ cells. n=4 independent LC donors. *p<0.05. D. Flow cytometry analysis of the percentage of CD4+IL10+ T cells after 5-day co-culture of migrated LC with and without dexamethasone stimulation, with CD4+ naïve T cells. 5-day cultures of CD4+ naïve T cells alone were used as control. n=5 independent LC donors. *p<0.05, **p<0.01.

### LC tolerogenic function depends on induction of IRF4-regulated transcriptional programme

We next sought to identify the mechanisms regulating transcriptional programming of immunotolerance in LCs. Single cell regulatory network inference analysis (SCENIC) in steady state and migrated LCs from the same skin donor identified 16 regulons in the steady state and 26 in migrated LCs (Aibar et al., 2017) (z-score enrichment >0.4, Figure 6A). In agreement with the observed induction of immunocompetence, regulons identified in migrated LCs were reported in immune cell activation (*JUND, STAT1, RELA, IRF4,* Figure 6A). To identify transcription factors important for the immune tolerance programming in LCs, 5 transcription factors with the highest changes in gene expression level were selected for partial information decomposition analysis in context (Figure 6B PIDC,^35^). PIDC was designed and benchmarked for GRN inference from single cell RNA-seq data, and is an extended formalism using multivariate information measures for each triplet of gene in the context of every cell in the dataset. However, since the information flow in a GRN flows from transcription factors to target genes, which expression transcription factors regulate, we decided to restrict direction of the edges within the inferred network, including only interaction edges consistent with the information flow (TF -> target gene, directional PIDC). The resulting network comprised 70 edges with weight higher than 1, and when hierarchically organised, predicted distinct regulatory modules for genes in the tol1 programme (Figure 6C, Supplementary Figure 6A-C). Interestingly, directional PIDC network analysis indicated combinatorial regulation of the majority of genes, with a single transcription factor implicated only for 3 targets. 6 target genes and 1 transcription factor were predicted to be regulated by *IRF4*. Analysis of our existing data from LCs with IRF4 expression edited by CRISPR-Cas9 confirmed, that expression of 3 out of the predicted genes was indeed compromised in IRF4 knock-down LCs (Figure 6D). Thus, LC migration from the epidermis results in a switching their transcriptional state resulting in the enhanced expression of a tolerogenic module that is dependent on *IRF4* regulation and underpins priming of Treg responses.

**Figure 6.**
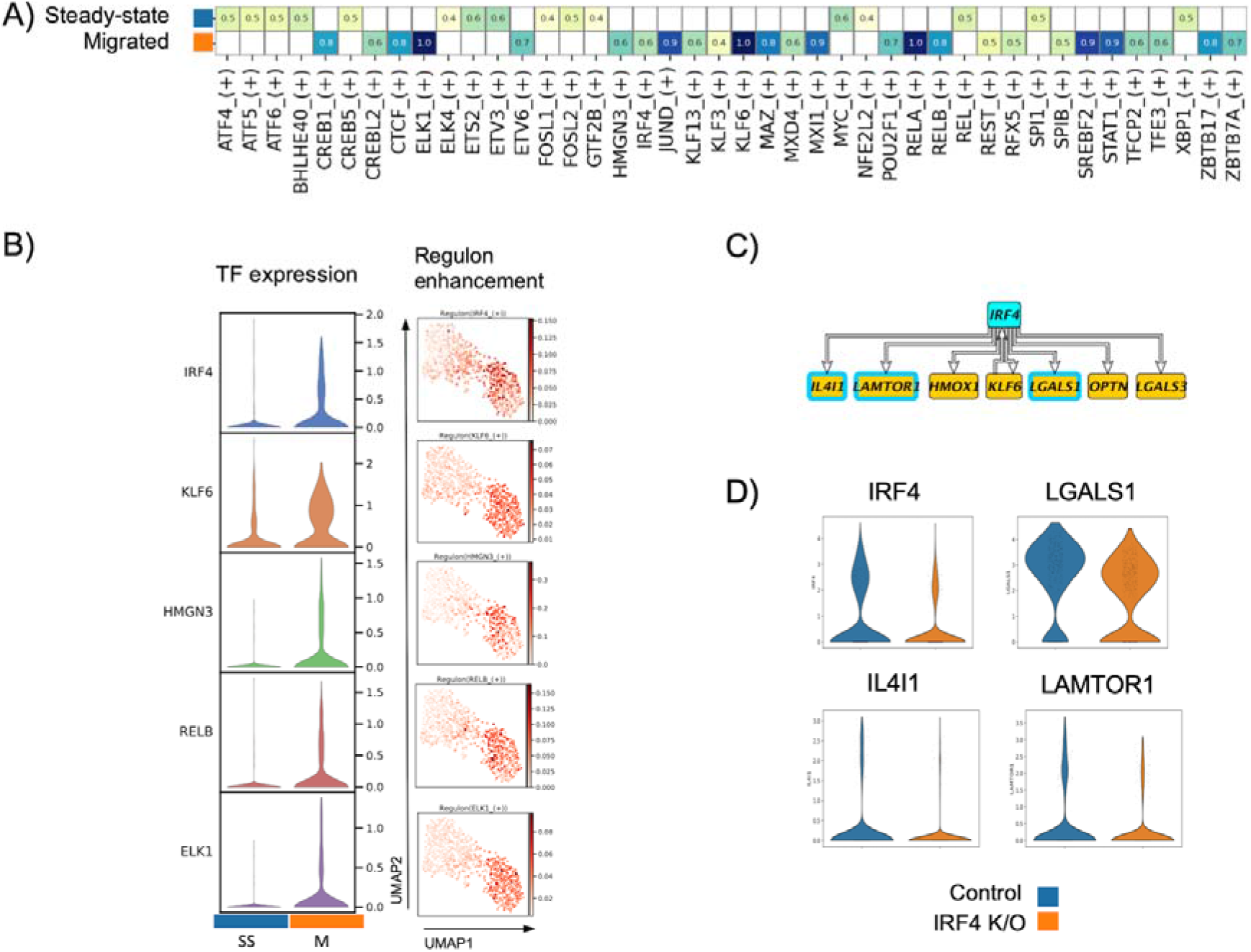
LC tolerogenic function depends on induction of IRF4-regulated transcriptional module. A. SCENIC regulatory network and inference clustering analysis revealed TF regulons which were enriched in steady-state and migrated LCs. Z-score heatmap of enriched regulons are displayed (z-score>0.4). B. Violin plots displaying the transcriptomic expression of TFs identified to be enriched in migrated LCs from SCENIC analysis. UMAP marker plots showing TF regulon enrichment Z-scores in each cell, across the two LC populations are displayed. C. Network displaying IRF4 with 6 target genes and 1 transcription factor as predicted by PIDC. D. 3 predicted IRF4 regulated genes (*IL4I1*, *LGALS1, LAMTOR1*) were identified to be downregulated in CRISPR-Cas9 IRF4 knock-down LCs.

## Discussion

While LC-mediated immunotolerance appears critical for cutaneous and systemic immune homeostatis, research into molecular mechanisms initiating and maintaining tolerogenic behaviour in human LCs has been significantly affected by the lack of appropriate experimental models and limitations of available technologies. Here, we applied a microfluidic-based single cell and single bead co-encapsulation (Drop-seq, ^22^) followed by high throughput sequencing of individual transcriptomes, to investigate transcriptional programmes in human LCs. This method allowed us to document heterogeneity in steady-state LC transcriptomes, displaying a spectrum of immune activation, previously, to the best of our knowledge, unobserved. Within this spectrum two subpopulations of steady-state LC could be identified: S1, characterised by low RNA content and immature state, and S2, which could be distinguished by high levels of CD86 expression at both mRNA and protein level, and displayed upregulated expression of antigen presenting genes, T cell co-stimulatory genes and genes generally associated with innate immune responses. Surprisingly, this latter population of LCs was characterised by ability to induce a tolerogenic T cell phenotype. Studies have previously shown that DCs in an immature and lowly activated state, expressing low levels of antigen presenting and co-stimulatory molecules, can drive tolerogenic responses by inducing anergy of antigen-specific T cells and expanding Tregs^36,37^. However, consistent with our findings, a study by Yamazaki et al. demonstrated that mature CD86High DCs were able to expand CD4+CD25+ T cells more effectively than CD86Low immature DCs^38^. Similarly, epicutaneous immunisation of mice increased migratory LC expression of CD80 and CD86 signature markers of activation, but did not result in efficient generation of effector memory CD4 T cells^39^. Our results indicate that while low expression levels of co-stimulatory molecules on immature DC reported by others may result in impaired generation of T cell activation, induction of tolerance requires delivering of efficient signal 2 through co-stimulation. Indeed, a study of transcriptional determinants of tolerogenic and immunogenic states in murine dendritic cells, highlighted that the DC antigen presentation gene module is overlaid by an IRF4-dependent regulatory programme in a tolerance-induction setting^40^. Supporting the association between immunocompetence and ability to induce tolerance, LC migration out of the epidermis resulted in an even greater ability to induce Tregs than immunocompetent (S2) steady-state LCs. Analysis of single LC transcriptomes confirmed that migratory LCs are mature and characterised by upregulation of MHC II molecules, co-stimulatory molecules (CD80 and CD86) and chemokine receptors (CCR7), indicating readiness for migration to local lymph nodes, mediating interaction with T cells to promote adaptive immune responses^41,42,43,19,32,33^.

The increase in T cell co-stimulatory genes, such as CD86 and MHC class II genes in the immunocompetent S2 cluster, and in migrated LCs, suggest that physical interaction with T cells is necessary for LCs to coordinate Treg induction. Indeed, CD86 activity itself has been implicated in DC-mediated tolerance induction through interaction with CTLA4 receptors on T cells^44,45^. However, since LC ability to induce tolerogenic responses is greatly enhanced with immunocompetence status upon migration, it is likely to be governed by additional factors complementing immune activation, the capacity to process and present antigen and interact with T cells via co-stimulatory molecules. Consistent with this possibility, we observed the presence of a tolerogenic gene module of 30 genes, including *IDO1*, *LAMTOR1, IL4L1* and *LGALS1,* expressed by LCs. Galectin-1 encoded by the *LGALS1* gene has been shown to promote the generation of tolerogenic DCs and to enable Tr1 type Tregs to supress Th1- and Th17-mediated inflammation^46,46^. Thus, Galectin-1 secreted by LCs could function in an autocrine as well as paracrine manner to promote Treg responses. The enzymes *IL4I1*, a mediator of H_2_O_2_ production and *HMOX1*, which degrades haem to carbon monoxide, have been shown to be expressed by DC and are implicated in the suppression of effector T cell activation and the induction of Tregs ^47–49^. Additionally, *LAMTOR1* is implicated in macrophage polarization towards an immunoregulatory M2 phenotype ^50^. Interestingly, PIDC analyses points to combinatorial regulation of the tolerogenic transcriptional programme in LCs. This is in concordance with the current understanding of gene transcription regulation, and highlights the importance of tolerance for LCs. While IRF4^51,52^, KLF6^53^ RELB^54,55^, and ELK1^56^ have been previously implicated in regulation of immunity or tolerance, *HMGN3* binds to nucleosomes and regultes chromatin organisation ^57^. Implication of its function in LCs are certainly intriguing, and warrant further detailed investigations. Importantly, analyses of IRF4 CRISPR-Cas9 knock-down in LCs confirm the dependence of several tolerance-related genes, including *IL4L1* and *LGALS1* and *LAMTOR1* on IRF4. Interestingly, IRF4 did not seem to regulate expression of IDO1, a classical tolerogenic mediator, which catabolises tryptophan leading to skewing of T cell differentiation towards Tregs^58^ directly. Identified as an interferon stimulated gene (ISG), IDO1 has been previously associated with DC activation by IFNs and TNF^59,60^ and has been implicated in a number of regulatory feedback loops in cross-talk with other cell types – e.g. activation of CTLA4 receptors on T cells in turn induces IDO1 expression in DCs^45^. Two studies in human LCs previously demonstrated induction of IDO1 steady-state LCs, and its importance for inhibition of effector T cell proliferation on stimulation with IFN-γ^61^ and for FcεRI signalling^62^. Our study confirms and extends these findings, highlighting IDO1 as a key regulator of LC tolerogenic responses induced during LC migration. Consistently, steady-state LCs, which do not express high levels of IDO1, have limited ability to activate regulatory T cells, are perhaps sufficient to maintain tolerogenic memory T cells *in situ*. In contrast, high levels of IDO1 induced by migration promote LC ability to prime naïve T cells for tolerance towards autoantigens.. Intrestingly, recent study documents, that IRF4 can form multipartite transcriptional complex with AHR, a well established inducer of IDO1 ^63,64^, binding to promoter elements of tolerance associated genes^65^. High levels of IRF4 expression would thus potentially promote more efficient AHR action, and increase in IDO1 expression. Induction of *AHR, IRF4* and *IDO1 axis* upon migration provides a mechanism for inducible IDO1 expression upon activation, and suggests the existence of a reinforcement loop for *in situ* tolerogenic responses through AHR-IRF4 signalling. This was previously observed in other DCs, where kynurenine metabolites, produced during IDO-mediated catabolism of tryptophan, feedback to AHRs to sustain IDO expression^66,63^.

Our extensive analysis of human primary steady-state and migrated LCs indicates that while LCs with tolerogenic ability exist *in situ*, migration greatly enhances LC tolerogenic potential. We postulate, that efficient priming of tolerogenic responses by LCs requires upregulation of a migration-coupled maturation program superimposed with a tolerance-inducing gene module. While the induction of this tolerogenic programme in LCs is complex, IRF4 is likely to act as a pivotal switch regulating LC immune function, and orchestrating complementary modules in LC transcriptional programming. The enhancement of LC tolerogenic abilities on maturation could be explored therapeutically to reinstate tolerance in the skin during inflammatory conditions.

## Acknowledgments

We are grateful to the subjects who participated in this study. We would like to thans Prof Peter Friedmann for in-depth review of the manuscript. We acknowledge the use of the IRIDIS High Performance Computing Facility and Flow Cytometry Core Facilities, together with support services at the University of Southampton. The study was funded by a Sir Hendy Dale Fellowship from Wellcome Trust, 109377/Z/15/Z. Development of single cell Drop-Seq technology was funded by MRC grant MC_PC_15078.

## Authorship Contributions

MEP, SS and JD: intellectually conceived and wrote the manuscript, planned the experiments and analysed the results SS, JD,KC,GP,AV: run functional experiments, flow cytometry, single-cell sequencing, AV, JD, PS, JW: developed and optimised scRNA-sequencing JD, BMA, AV, PS, MEP: analysis and meta-analysis of scRNA-seq data MEP, MAJ, HS: discussions, data analysis, reviewing of the manuscript

## Supplementary Material

**Supplementary Table 1. Trypsinised steady-state LC and TolMoDC DEGs**

DEGs comparing trypsined steady-state LC to unstimulated MoDC (GSE23618) and TolMoDC to unstimulated MoDC (GSE52894) were indentified using Limma (FDR corrected p-value<0.05, logFC>1). Biological pathways associated with DEGs were identified in Toppgene (FDR corrected p-value<0.05).

**Supplementary Table 2. Steady state LC DEG analysis**

DEGs comparing steady-state S1 and S2 LC using Limma (FDR corrected p-value<0.05, logFC>1). Biological pathways associated with DEGs were identified in Toppgene (FDR corrected p-value<0.05).

**Supplementary Table 4. Tolerogenic DC gene signature 1.**

Literature reviews and experimental papers referencing genes associated with DC and macrophage tolerogenic function were summarised into a 64 gene signature.

**Supplementary Table 3. Migrated and Steady state LC DEG analysis**

DEGs comparing migrated and steady-state LC using Limma (FDR corrected p-value<0.05, logFC>1). Biological pathways associated with DEGs were identified in Toppgene (FDR corrected p-value<0.05).

## Supplementary Figures

**Figure S1.**
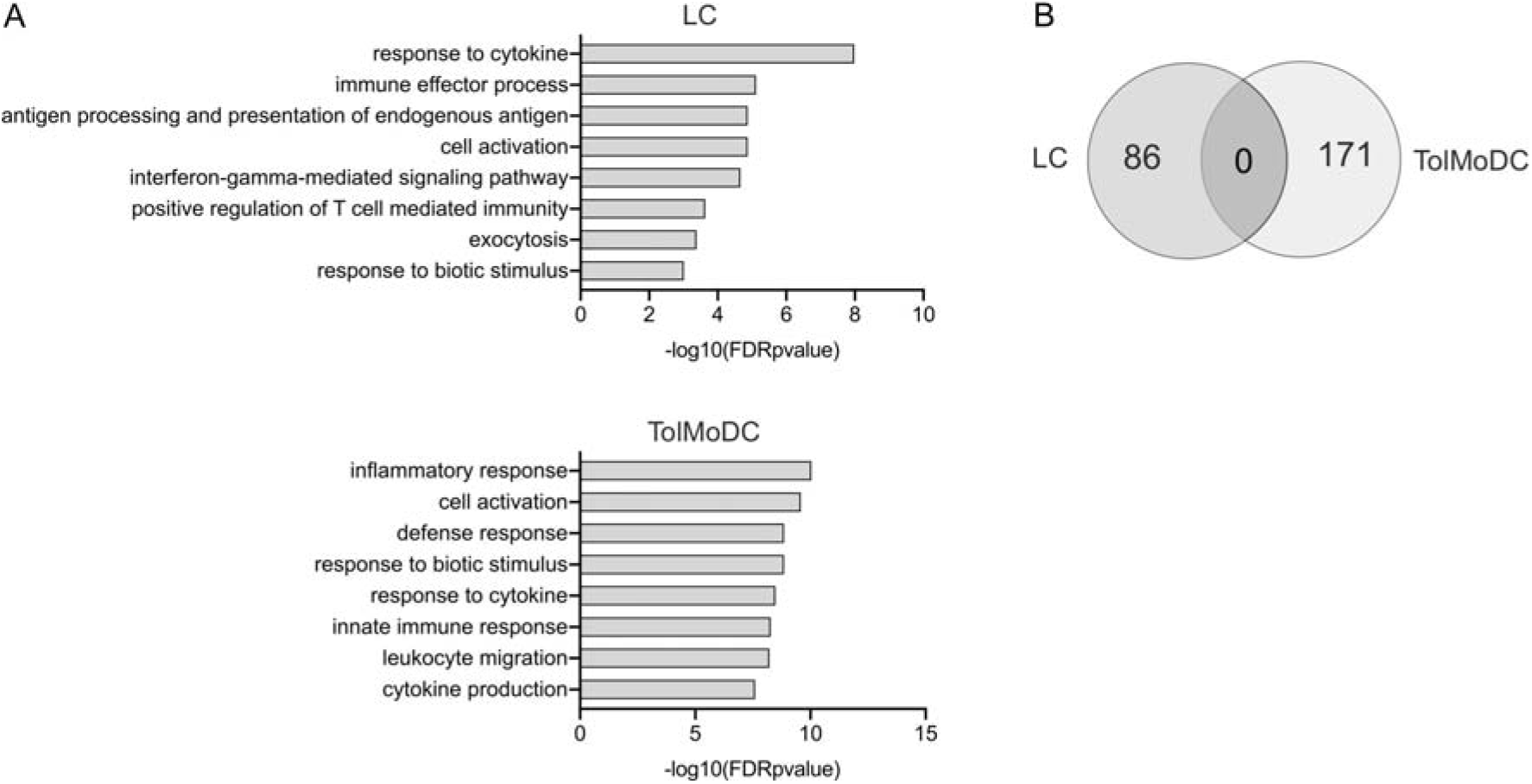
Steady state LCs exist in a spectrum of immune activation from immaturity to immunocompetency. A. Biological pathways identified from gene ontology analysis (ToppGene) of 86s DEGs upregulated in trypsinised steady-state LC compared unstimulated MoDC (GSE23618) and 171 DEGs upregulated in Dexamethasone and Vitamin D3 treated MoDC (TolMoDC) compared to unstimulated MoDC (GSE52894) identified using Limma (FDR corrected p-value<0.05, logFC>1). -log(10) FDR corrected p-values are displayed. B. Venn diagram displaying crossover between upregulated DEGs identified comparing in trypsinised steady-state LC to unstimulated MoDC (GSE23618), and TolMoDC to unstimulated MoDC (GSE52894).

**Figure S2.**
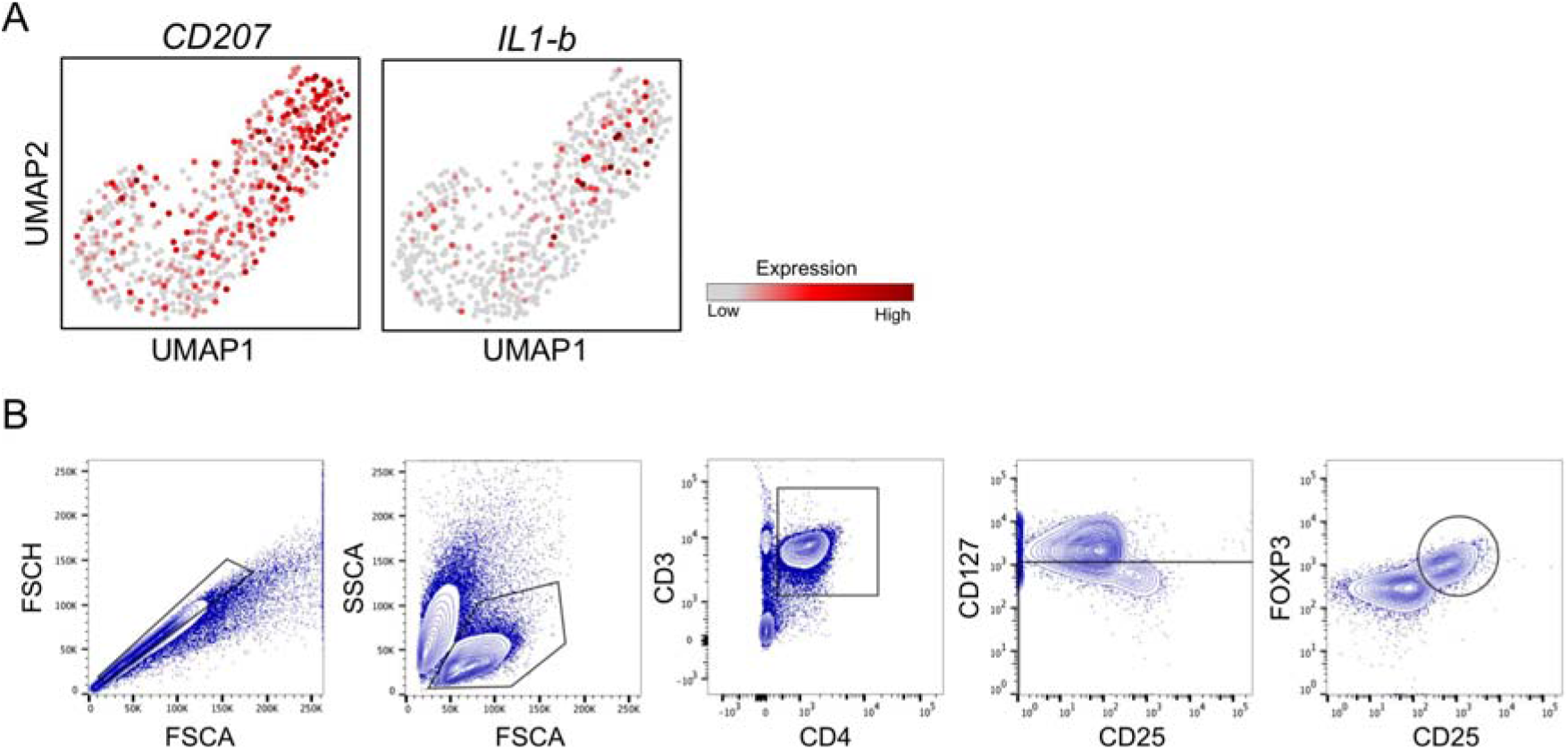
Steady state immunocompetent LCs are superior at inducing FOXP3+ Tregs. A. UMAP markers plots displaying *CD207* and *IL1B* expression amongst the steady-state LC population displaying low (grey) to high (dark red) SCnorm normalised expression. B. Gating strategy for investigating the quantity of CD3+CD4+CD127-CD25+FOXP3+ Tregs after co-culture of CD4 naïve T cells with LC for 5-days.

**Figure S3.**
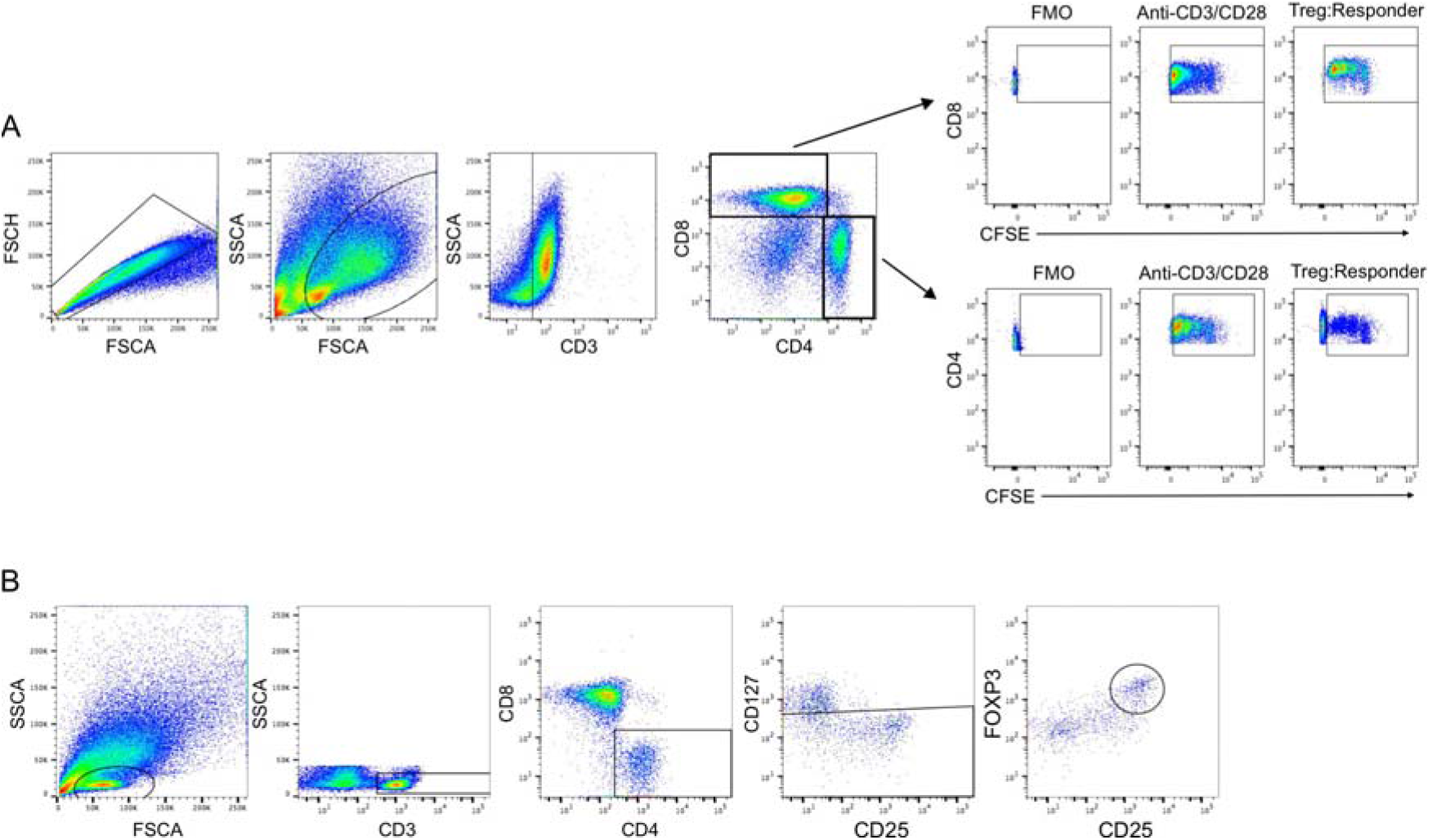
Migration enhances tolerogenic abilities of immunocompetent LCs. A. Gating strategy for investigating CFSE labelled PBMC proliferation, selecting for CD4+ and CD8+ T cell populations. CFSE measurement gating was applied to responder cell population only, excluding unlabelled CFSE negative Tregs. B. Gating strategy for investigating the quantity of Tregs induced after co-culture of autologous TRMs with migrated LC for 5-days.

**Figure S4.**
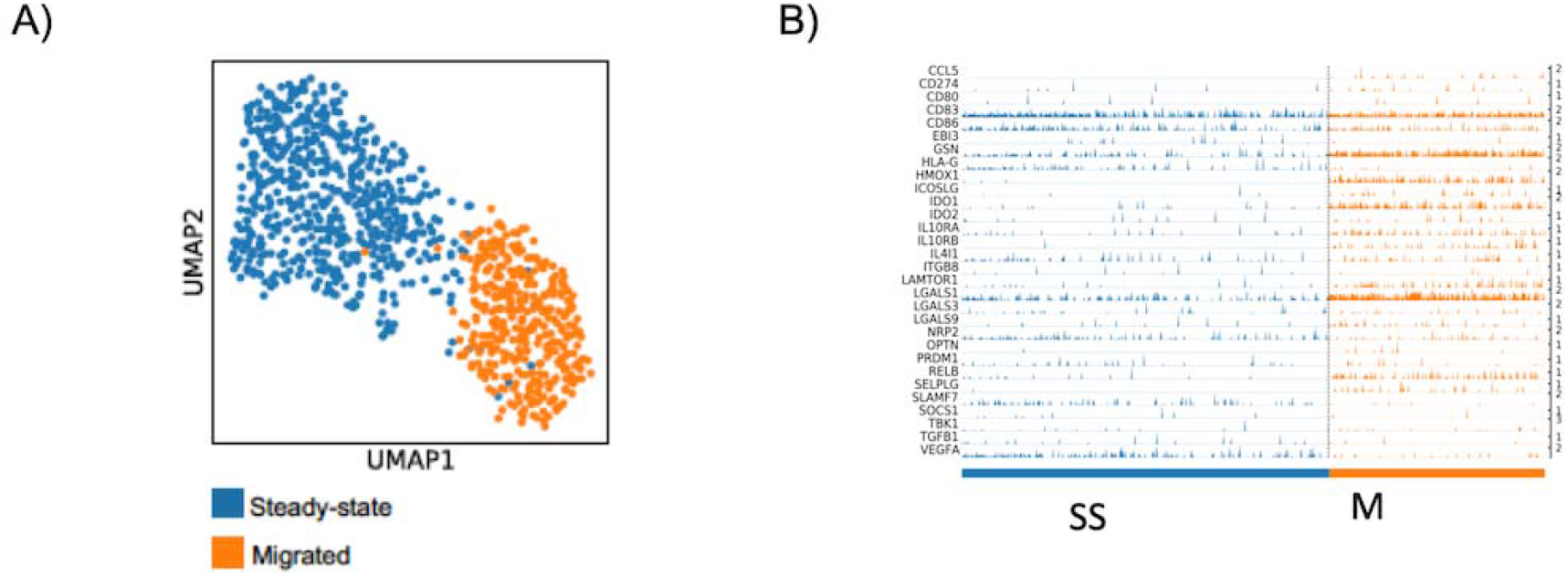
Migration from the epidermis enhances LCs tolerogenic programming. **A.** UMAP markers plots displaying steady state (blue) and migrated (orange) LC. SCnorm normalised expression. **B.** Trackplot of all 30 genes from the tol 1 signature which are within the whole LC single cell transcriptomic dataset.

**Figure S5.**
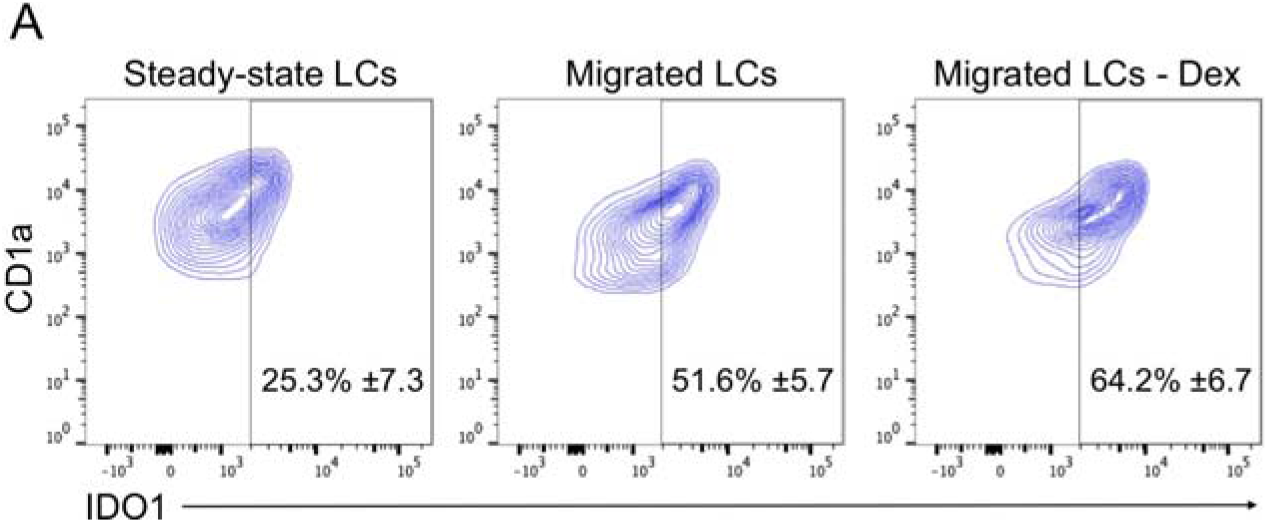
LC-induced tolerance is mediated by IDO1 can be enhanced by immunotherapeutic intervention. A. Gating strategy to define the percentage of IDO1 expression in steady-state LC, migrated, and dexamethasone migrated LC. n=5 steady-state and migrated LC experiments, n=4 migrated dexamethasone experiments.

**Figure S6.**
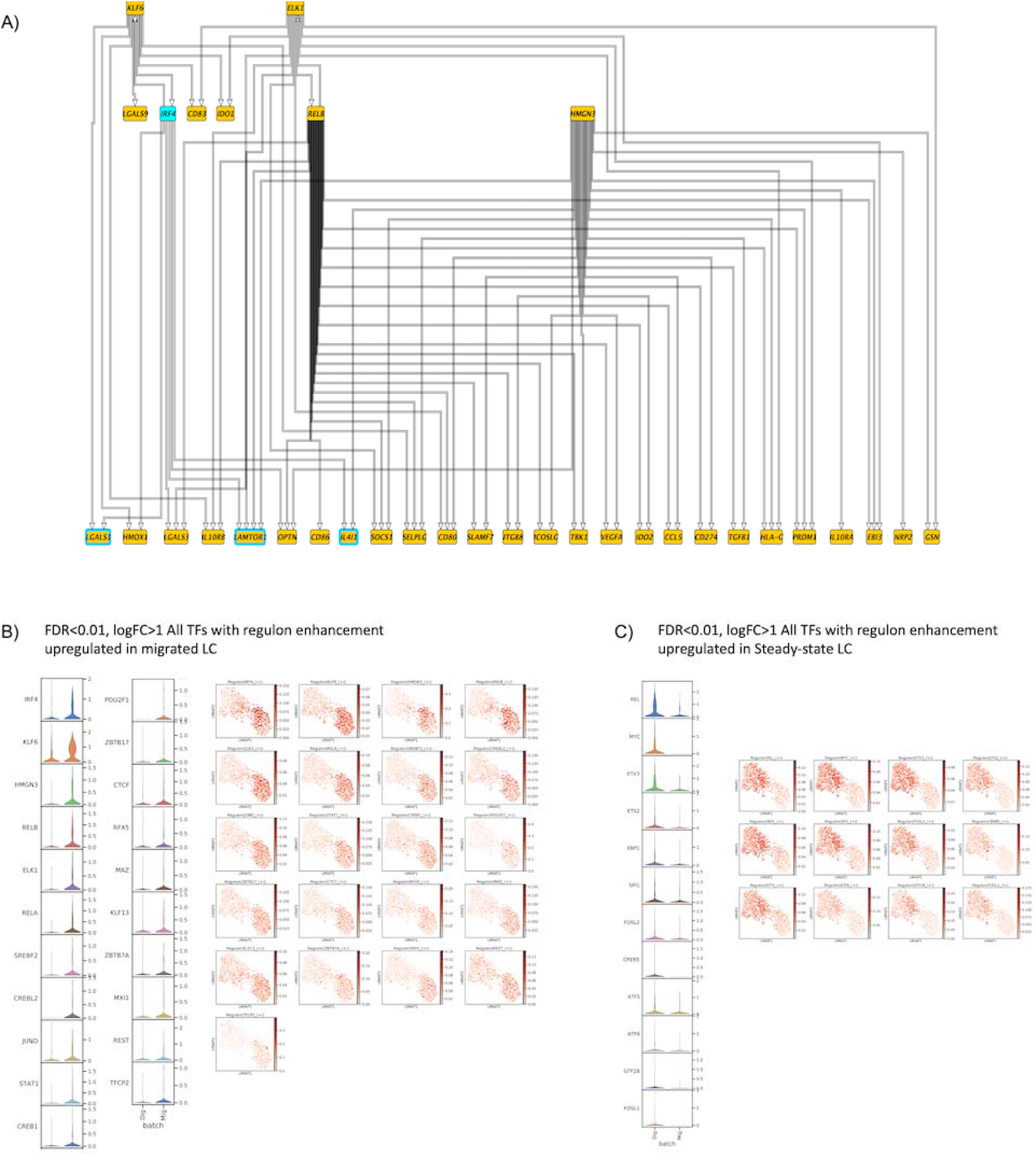
LC tolerogenic function depends on induction of IRF4-regulated transcriptional module. **A.** PIDC network graph comprising 70 edges with weight >1, hierarchically organized, displaying predicted regulatory modules for the top 5 enriched TFs with genes within the tol 1 signature. **B.** Violin plots displaying the transcriptomic expression of migrated LC upregulated TFs (FDR corrected p-values<0.01, logFC>1) identified to be enriched in migrated LCs from SCENIC analysis (z-score>0.4). UMAP marker plots showing TF regulon enrichment Z-scores in each cell, across the two LC populations are displayed alongside. **C.** Violin plots displaying the transcriptomic expression of steady-state LC upregulated TFs (FDR corrected p-values<0.01, logFC>1) identified to be enriched in steady-state LCs from SCENIC analysis (z-score>0.4). UMAP marker plots showing TF regulon enrichment Z-scores in each cell, across the steady-state and migrated LC populations and datasets (D1-D4) are displayed alongside.

